# Essential Molecular Variables of liverworts in biological soil crusts

**DOI:** 10.64898/2026.07.16.738858

**Authors:** Kristian Peters, Nicole M. Van Dam, Steffen Neumann

## Abstract

(1) In plant ecology, functional traits are widely used to predict plant performance in ecosystems. Little is known on molecular traits and how they integrate with the classical trait spectrum, in particular in non-vascular land plants. We conceptualize Essential Molecular Variables (EMVs) using complex-thallose liverworts in biocrusts as reference.

(2) We combined morphometry and bioimaging with liquid chromatography high-resolution mass-spectrometry (UPLC/ESI-QTOF-MS) with data-dependent acquisition of tandem mass-spectra (DDA-MS). To identify EMVs, data mining was performed including regression trees, random forest and redundancy analysis combined with manual filtering and functional annotation using literature and chemical libraries.

(3) We found the molecular trait spectrum to be highly structured and largely orthogonal to classical traits. We identified six major categories of EMVs: molecule structure, carbon metabolism, amino acids and peptides, shikimates and phenylpropanoids, terpenoids and alkaloids that were functionally related to biotic interactions and bioclimatic properties.

(4) We find EMVs to be indicators that characterize plant performance in ecosystems. Molecular traits and EMVs represent the mechanistic link to form and function and extend the functional trait concept. They provide mechanistic insights into how global changes destabilize liverworts in biocrusts and represent a new tool set by informing functional ecology on molecular mechanisms.

## Introduction

In plant ecology, functional traits are widely used to predict plant performance in ecosystems and ecosystem functioning (Bruelheide *et al*., 2018). The diversity of plant forms and life-histories is largely explained by two components, whereby the first corresponds to plant size-related traits and the second relates to the leaf economics spectrum, which balances leaf construction costs with growth, survival and reproduction (Wright *et al*., 2004; Díaz *et al*., 2016). Whereas the mechanisms of classical traits, that is morphological, physiological and phenological traits, of vascular plants are comparably well understood, still little is known on molecular traits (Walker *et al*., 2023). They have been shown to provide additional dimensions to the spectrum of form and function (Walker *et al*., 2022) and yield insight into how the metabolome varies across the plant kingdom while also providing mechanistic insights (Walker *et al*., 2023). Molecular traits can be represented by all kinds of molecules, including metabolic compounds, proteins, or transcribed RNA. Here, we focus on metabolic compounds, molecular descriptors and molecular families. The latter comprise compound classes, e.g. natural product classes (NPC) such as terpenoids or flavonoids which aggregate metabolites according to biosynthetic pathways (Kim *et al*., 2021b). In addition, we consider molecular descriptors, which describe physicochemical properties of molecules such as polarity, mass, or molecular complexity (Walker *et al*., 2023).

The assessment of molecular traits results in a highly multi-dimensional matrix often accompanied with risk of overfitting statistical models. Therefore, a simplification may offer a more abstracted and ecologically meaningful look at mechanistic processes (Carmona *et al*., 2016). To reduce the dimensionality of the molecular trait matrix, we here hypothesize a minimum set of Essential Molecular Variables (EMVs) that contain all essential information to predict plant performance in ecosystems and, eventually, ecosystem functioning. EMVs summarize molecular traits or groups of linked molecular variables to only a few robust aggregated variables which can be used to extrapolate their functioning at coarser ecological scales. The resulting EMVs should serve as indicators that abstract low-level molecular observations to be indicative for higher-level ecological functions. By generalizing molecular traits into categories of EMVs, we follow the same principle as with defining essential biodiversity variables (EBVs) for biodiversity (Pereira *et al*., 2013), but now aiming to bridge molecular and functional ecology. By providing detailed molecular mechanisms that shape plant function through evolutionary and ecological processes, our novel EMV-concept also helps to advance the functional trait concept itself (Violle *et al*., 2007).

While first steps have been made to integrate molecular and classical traits in explaining plant form and function (Walker *et al*., 2023), we have no information on the role of molecular traits in non-vascular plants and especially in predicting plant performance in ecosystems. Important ecosystems that are dominated by non- vascular plants are biological soil crusts, which are an ensemble of microscopic (cyanobacteria, algae, fungi, bacteria, and protists) and macroscopic (lichens, bryophytes, and microarthropods) poikilohydric organisms that occur on or within the top few centimeters of the soil surface (Navas Romero *et al*., 2020; Weber *et al*., 2022). Complex-thallose liverworts dominate these biocrust communities. They form mutualisms with filamentous cyanobacteria, soil-bound bacteria such as actinobacteria or betaproteobacteria that exude polysaccharide sheaths and bind soil particles to allow for subsequent establishment (Steel *et al*., 2004; Seppelt *et al*., 2016; Cheng *et al*., 2024). To test and to identify EMVs, liverworts particularly qualify as good study subjects as they have a comparably basic biology, a short life-span, a sudden response to biotic and abiotic changes, and close evolutionary relationship to the model species *Marchantia polymorpha* (Peters *et al*., 2022). In addition, trait- habitat interactions of bryophytes in biocrusts are comparably simple, which facilitates associating molecular traits to functions at coarser ecological scales (Seppelt *et al*., 2016; Coe *et al*., 2019).

Despite research on liverworts within biocrusts, we still have a poor understanding of functional trait relationships of liverworts and their impact on biodiversity and ecosystem functioning (Stanton & Coe, 2021; Van Zuijlen *et al*., 2023). In contrast to vascular plants, leaves of many bryophyte species consist only of a single layer of poikilohydric cells where investment in structure is minimal. On the other side of the spectrum, complex-thallose liverworts do not produce leaves and form a complex thallus that represents the entire plant including photosynthetically active tissue. The spectrum of form and function and whether the importance of leaf-based traits like specific leaf area or leaf economics are similarly important in liverworts as in vascular plants is much less investigated (Coe *et al*., 2024). To better understand the functional role of the trait spectrum and to conceptualize EMVs, we investigate phenotypic and molecular traits of 14 non-vascular bryophyte species of complex thallose liverworts (order Marchantiales) in 16 biocrusts (Fig. 1). Phenotypic traits were acquired using morphometry and bioimage analysis, and molecular traits using liquid chromatography high-resolution mass-spectrometry (UPLC/ESI-QTOF-MS) with data-dependent acquisition of metabolite-characterizing tandem mass-spectra (DDA-MS), respectively.

**Figure 1.**
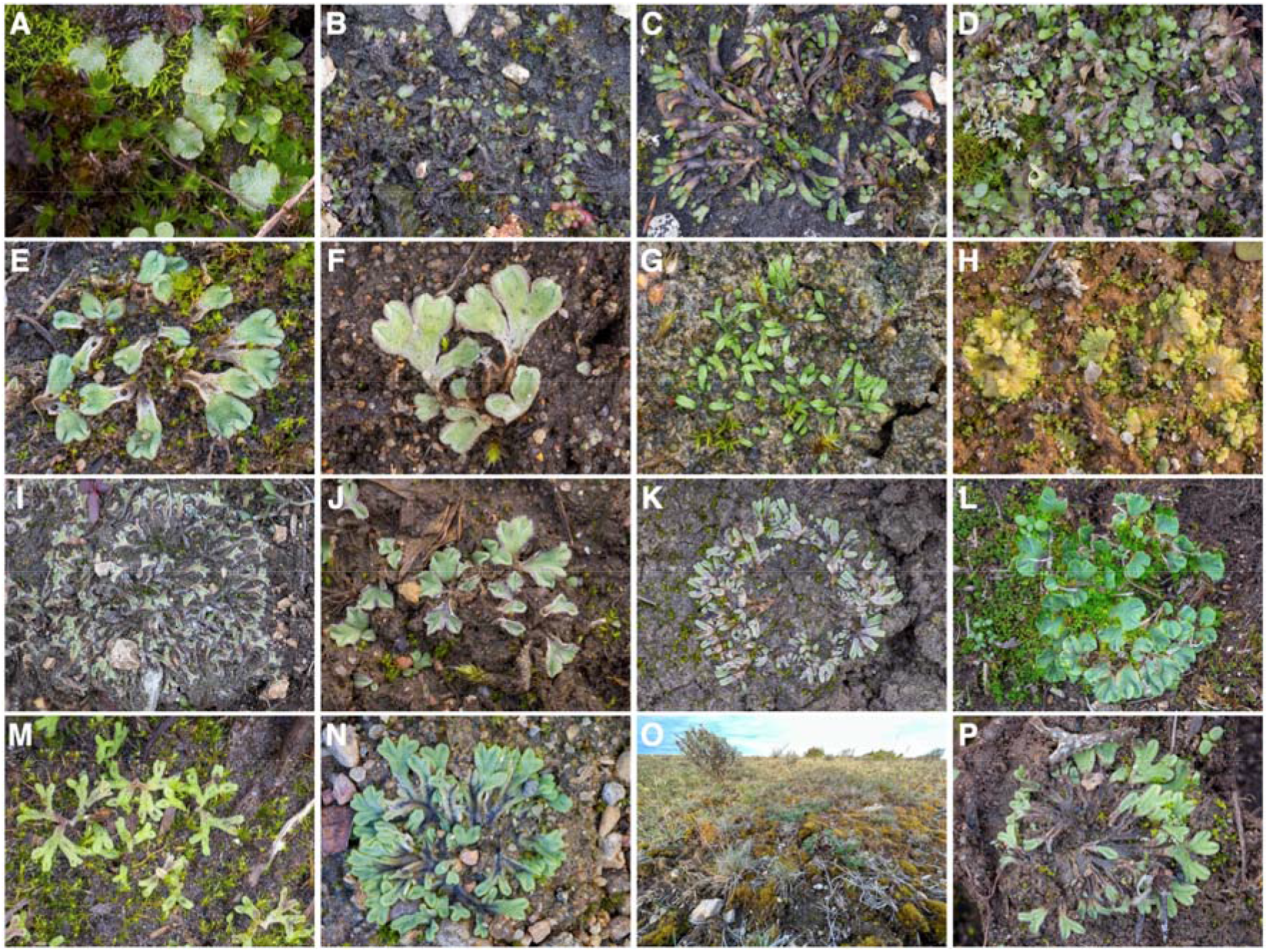
Stature images and overview on the investigated species. **(a)** *Asterella gracilis* (F.Weber) Underw. **(b)** *Athalamia hyalina* var. *suecica* (Lindb. ex Gottsche & Rabenh.) S.Hatt. **(c)** *Mannia fragrans* (Balb.) Frye & L.Clark **(d)** *Reboulia hemisphaerica* subsp. *hemisphaerica* (L.) Raddi **(e)** *Riccia beyrichiana* Hampe **(f)** *Riccia bifurca* Hoffm. **(g)** *Riccia canaliculata* Hoffm. **(h)** *Riccia cavernosa* Hoffm. **(i)** *Riccia ciliifera* Link GER1 **(j)** *Riccia ciliifera* Link SWE **(k)** *Riccia gothica* Damsh. & Hallingb. **(l)** *Riccia ciliifera* Link GER2 **(m)** *Riccia huebeneriana* Lindenb. **(n)** *Riccia sorocarpa* Bisch. **(o)** *Riccia subbifurca* Warnst. ex Croz. SWE1 **(p)** *Riccia subbifurca* Warnst. ex Croz. SWE2.

Using liverworts as reference, the aims of this study are (1) to integrate the classical and molecular trait spectrum of the chosen liverworts and compare them to those of vascular plants, (2) to structure the molecular trait spectrum to potentially reveal EMVs as molecular indicators of plant performance in ecosystems, and (3) to make conclusions on the liverwort’s performance in biocrusts. We hypothesize that the liverworts evolved general and species-specific phenotypic and molecular mechanisms that do not appear to be random but to be functionally grouped. These molecular traits link plant performance to different conditions in biocrusts and extend the classical trait spectrum. We further hypothesize that the molecular spectrum of form and function is structured allowing for defining major categories of EMVs.

## Materials and Methods

### Sampling

Samples were collected in the field and put immediately in sterile petri dishes at locations in Southern Sweden in September 2022 and in Germany in October 2022 (Table 1). Specimens were photographed on-site and bioimaging was performed on fresh material using macro- and microscopy in the lab. After an incubation period of five days at room temperatures, the plant material was isolated, washed under a light microscope to remove dirt and other residues, filled into Eppendorf tubes and shock-frozen (LC-MS analyses, three biological replicates per sample), or dried (marker sequencing, one replicate per sample). Remaining plant material was stored as voucher specimens in the Herbarium Haussknecht Jena (barcodes: JE04010739- JE04010754) (Table 1).

**Table 1:** List of voucher specimens and fresh samples investigated in this study. The columns list the taxonomic species identifiers (*NCBI*, *GBIF*), the family the sample belongs to within the order of Marchantiales, geographic coordinates (WGS84 format, all coordinates have ± 10 m accuracy), sampling data and the voucher specimen identifiers of the Herbarium Haussknecht Jena. The first letters indicate the *Index Herbariorum* institution code (Holmgren & Holmgren, 1991).

| Sample ID | Species names | Family | Taxonomy IDs | Geographic coordinates | Sampling date | Voucher identifiers |
| --- | --- | --- | --- | --- | --- | --- |
| **Asterella gracilis SWE** | *Asterella gracilis* (F.Weber) Underw. | Aytoniaceae | NCBI:122620<br>GBIF:5286285 | 58.030607002,<br>16.455699317 | 2022-09-16 | JE4010742 |
| **Athalamia hyalina SWE** | *Athalamia hyalina* var. *suecica* (Lindb.<br>ex Gottsche & Rabenh.) S.Hatt. | Cleveaceae | NCBI:122615<br>GBIF:12364697 | 57.81875188,<br>18.573477554 | 2022-09-17 | JE4010741 |
| **Mannia fragrans SWE** | *Mannia fragrans* (Balb.) Frye & L.Clark | Aytoniaceae | NCBI:179045<br>GBIF:5286289 | 57.81875188,<br>18.573477554 | 2022-09-17 | JE4010739 |
| **Reboulia hemisphaerica SWE** | *Reboulia hemisphaerica* subsp.<br>*hemisphaerica* (L.) Raddi | Aytoniaceae | NCBI:37395<br>GBIF:7679096 | 57.81875188,<br>18.573477554 | 2022-09-17 | JE4010740 |
| **Riccia beyrichiana SWE** | *Riccia beyrichiana* Hampe | Ricciaceae | NCBI:122635,<br>GBIF:8550835 | 56.588578513,<br>16.60593546 | 2022-09-19 | JE4010749 |
| **Riccia bifurca SWE** | *Riccia bifurca* Hoffm. | Ricciaceae | NCBI:2779757<br>GBIF:5710080 | 57.848309241,<br>18.633133946 | 2022-09-17 | JE4010752 |
| **Riccia canaliculata SWE** | *Riccia canaliculata* Hoffm. | Ricciaceae | NCBI:2779758<br>GBIF:5710075 | 57.79400221,<br>18.484261361 | 2022-09-17 | JE4010747 |
| **Riccia cavernosa SWE** | *Riccia cavernosa* Hoffm. | Ricciaceae | NCBI:122636<br>GBIF:5710073 | 55.935140445,<br>14.261055376 | 2022-09-15 | JE4010743 |
| **Riccia ciliifera GER1** | *Riccia ciliifera* Link | Ricciaceae | NCBI:71659<br>GBIF:9053960 | 51.503291273,<br>11.94492469 | 2022-10-13 | JE4010753 |
| **Riccia ciliifera GER2** | *Riccia ciliifera* Link | Ricciaceae | NCBI:122638<br>GBIF:9362069 | 51.503291273,<br>11.94492469 | 2022-10-13 | JE4010754 |
| **Riccia ciliifera SWE** | *Riccia ciliifera* Link | Ricciaceae | NCBI:71659<br>GBIF:9053960 | 57.845770321,<br>18.674179874 | 2022-09-17 | JE4010748 |
| **Riccia gothica SWE** | *Riccia gothica* Damsh. & Hallingb. | Ricciaceae | GBIF:7456038 | 57.820616523,<br>16.571632151 | 2022-09-17 | JE4010746 |
| **Riccia huebeneriana SWE** | *Riccia huebeneriana* Lindenb. | Ricciaceae | NCBI:122639<br>GBIF:5836563 | 56.361714341,<br>13.850084042 | 2022-09-15 | JE4010745 |
| **Riccia sorocarpa SWE** | *Riccia sorocarpa* Bisch. | Ricciaceae | NCBI:122646<br>GBIF:5286296 | 56.358981715,<br>14.541635529 | 2022-09-15 | JE4010744 |
| **Riccia subbifurca SWE1** | *Riccia subbifurca* Warnst. ex Croz. | Ricciaceae | NCBI:1914413<br>GBIF:6096600 | 56.528407722,<br>16.521946862 | 2022-09-19 | JE4010750 |
| **Riccia subbifurca SWE2** | *Riccia subbifurca* Warnst. ex Croz. | Ricciaceae | NCBI:1914413<br>GBIF:6096600 | 56.588578513,<br>16.60593546 | 2022-09-19 | JE4010751 |

### Microscopy, image processing and bioimage analysis

Microscopy and acquisition of raw bioimages was conducted using the methods described in (Peters & König-Ries, 2022).

Morphometric characters were measured three times on three biological replicates per sample and were performed with Fiji (ImageJ version 2.14.0/1.54f) (Schindelin *et al*., 2012). After setting up the scales for the individual images in the Image- Properties menu, the following morphometric characters were measured manually: thallus width [µm], thallus length [µm], thallus with violet pigments [0/1], ventral scales [0/1], ventral scales with slime cells [0/1], ventral scales with violet pigments [0/1], ventral scales with hairs [0/1], air pores [0/1], width of ring cells of air pores in adaxial view [µm], height of ring cells of air pores in cross section [µm], number of ring cells of air pores in cross section [#], width of ring cells of air pores in cross section [µm], height of ring cells of air pores in cross section [µm], width of epidermis cells in cross section [µm], height of epidermis cells in cross section [µm], width of subepidermal cells in cross section [µm], height of subepidermal cells in cross section [µm], width of thallus in cross section [µm], height of thallus in cross section [µm], height of thallus wing in cross section [µm], angle of thallus wing in cross section [°], width of thallus wing in cross section [µm], area of thallus in cross section [µm^2^] (Figure 1, Supplemental Table S1). The protocol of (Waite & Sack, 2010) served as a reference and was extended to complex-thallose liverworts, accordingly. Lengths and widths were obtained using the Measure function from the Analyze menu and the results saved in CSV files. To automate the measurement of area, a pixel classification model was trained using LabKit (Arzt *et al*., 2022) by selecting representative foreground and background areas. The imaging data was segmented subsequently with StarDist (Weigert *et al*., 2020) using the above classifier. StarDist automatically classified objects in the images into foreground and background and areas classified as foreground were then measured using the Measure function from the Analyze menu and results were saved. CSV files with all individual morphometric measurements of all specimens were joined into one single table and used for subsequent data analyses.

In order to represent the high plasticity and variability of morphological characters of liverworts, variance, skewness and kurtosis of any of the measured continuous morphometric characteristics were calculated additionally according to (Enquist *et al*., 2015; Zhang *et al*., 2023).

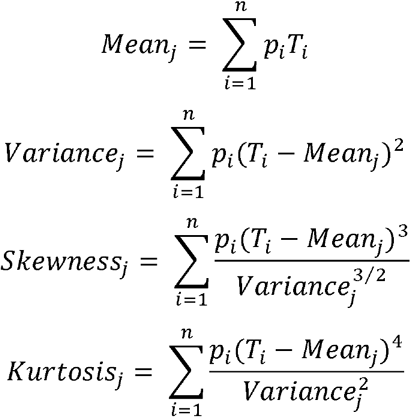

For each character, *p_i_* represents the measurements for species *j*, *T_i_* the mean value of character measurements of all species, *n* is the total number of measurements for the character. The variance and skewness represent the dispersion range and asymmetry of the character distribution, respectively. The kurtosis indicates the relative peak of the character distribution and the heaviness of its tails. Low kurtosis reflects high evenness in abundance of character values which implies similarity of species.

Raw camera and pre-processed imaging data in CR3 and TIFF format, respectively, were deposited to the BioImage Archive (BioStudies) using the command line IBM Aspera software tool ascp version 3.8.1.161274. Processed images and the metadata were deposited to the Image Data Resource.

### DNA marker sequencing

DNA marker sequences of the *trn*L-*trn*F (*trn*LF) plastid region were obtained by the German Barcode of Life using liverwort-specific primers described in (Stech & Quandt, 2014). Sequences were analyzed using an AB 3730 DNA analyzer instrument and PeakTrace was used as basecaller and trace processor. Multiple sequences were aligned using the MUSCLE 3.8 REST web service Perl client (Edgar, 2004). The resulting FASTA file was used for further phylogenetic analyses. Sequencing data was deposited to the European Nucleotide Archive (ENA) (Burgin *et al*., 2023) and is available under the study identifier ERP155252 (accession PRJEB70317) (https://www.ebi.ac.uk/ena/browser/view/PRJEB70317). Raw reads of *trn*LF sequences of the 16 samples are available under the sample identifiers SAMEA114863468-SAMEA114863483.

### Untargeted LC-MS-MS Analysis

We followed extraction procedures for LC-MS originally developed for vascular plants by (Böttcher *et al*., 2009) and adapted for the use with bryophytes and the different analytical setup used herein by (Blatt-Janmaat *et al*., 2022). This data-dependent acquisition method has been shown to provide robust results and high-quality spectra for thallose liverworts (Peters *et al*., 2023). In detail, frozen plants were homogenized for 5 minutes at 30 Hz in Teflon chambers equipped with two stainless steel balls. The chambers were rinsed with 100 µL of cold 80:20 MeOH:H2O supplemented with 2 µM 4-Methylumbelliferon, 5µM Kinetin (Sigma), 5µM Biochanin A (Sigma), and 5 µM N-(3-Indolylacetyl)-L-alanine (Sigma) and the slurry was shaken at 1000 rpm for 15 minutes at room temperature. The mixture was centrifuged twice at 13000 rpm for 15 minutes and the supernatant was transferred to autosampler vials.

Samples were separated with an Agilent 1290 Infinity HPLC (Agilent, Waldbronn, Germany) equipped with a Nucleodur X18 Gravity-SB column (1.8µm 100x2 Macherey Nagel, Dueren, Germany) coupled to a Bruker TIMS-TOF-MS (timsTOF Pro, Bruker, Bremen, Germany). Separations were performed at 35°C with the following binary gradient of 0.1% aqueous formic acid (solvent A) and acetonitrile (solvent B): isocratic 98% A for 1 minute; a linear gradient from 98% to 5% A from 1 to 14 minutes; isocratic 5% A from 14 to 17.5 minutes; isocratic 98% A from 18 to 20 minutes. The flow rate was maintained at 0.5mL/min, the injection chamber was maintained at 4°C. Injections were performed separately for positive and negative ionization mode. Data-dependent acquisition (DDA-MS) mode was used with the instrument settings described in (Blatt-Janmaat *et al*., 2022).

Raw data converted into mzML format using msconvert (Chambers *et al*., 2012) as well as derived data (SIRIUS project folders, RData) were deposited in MetaboLights under the study identifier MTBLS2239 (Haug *et al*., 2013). Metadata were recorded in compliance with the minimum information guidelines for Metabolomics studies (Spicer *et al*., 2017).

Data processing was performed in the statistical software environment R version 4.3.1 using the iESTIMATE framework (https://github.com/ipb-halle/iESTIMATE).

Chromatographic peak detection was performed using the R package XCMS version 3.22.0 (Smith *et al*., 2006). The following settings were used for the negative ion mode: CentWaveParam, ppm = 25, mzCenterFun = “mean”, peakwidth = c(7.2, 36), prefilter = c(2, 50), mzdiff = 0.0012, snthresh = 7, noise = 0, integrate = 1, firstBaselineCheck = TRUE, verboseColumns = FALSE, fitgauss = FALSE, roiList = list(), roiScales = numeric()). For the positive ion mode different settings were used for: peakwidth = c(9.4, 32), prefilter = c(6, 51), mzdiff = -0.0043, snthresh = 2. Grouping of chromatographic peaks was performed before and after retention time correction with the following settings in both ion modes: PeakDensityParam, minFraction = 0.7, bw = 0.25, minSamples = 1, binSize = 0.5. Retention time correction was performed with the following settings: PeakGroupsParam, minFraction = 0.7, smooth = “loess”, span = 0.2, family = “gaussian”. Only metabolite features with retention times less than 1020 s were considered for further analysis.

The MS1-level peak tables were created separately for positive and negative ion modes with the settings featureValues, method = “medret”, value = “into”. The peak tables were log2-transformed, and missing values were imputed with zeros.

Histograms and PCA diagnostic plots were generated to additionally evaluate the distribution of the data. MS2-level fragment spectra (MS-MS spectra) that were acquired by the Data-Dependent Acquisition mode (DDA-MS) were extracted from the profiles using the chromPeakSpectra, msLevel = 2 L, return.type = “Spectra” settings of XCMS. Spectra obtained from the same precursor ion were combined using the combineSpectra function from the R package Spectra using the following settings: FUN = combinePeaks, ppm = 25, peaks = “union”, minProp = 0.8, intensityFun = median, mzFun = median, backend = MsBackendDataFrame. These steps were performed separately for positive and negative ion modes. The MS1-level peak tables were then filtered to include only peaks for which the DDA-MS had acquired MS-MS fragment spectra. The spectra were saved in MSP and MGF files for further data processing. The peak tables and associated spectral and annotation data for positive and negative modes have been made separately available in MetaboLights.

As standard variance and median values deviated less than 5%, the filtered MS1- level peak tables containing log-transformed abundances of peaks in positive and negative ion modes were joined and used for further statistical analyses.

Presence/absence peak tables were also generated to contain whether a metabolite feature was detected in the profiles. Features with abundances less than 10^−8^ % of the median abundance were considered not detected.

Annotation and classification of MS-MS fragment spectra was carried out with the software SIRIUS version 5.7 (Dührkop *et al*., 2019a). The settings described in (Peters *et al*., 2023) were used for both ionizations. Annotation was accomplished automatically by selecting the highest-ranking candidate for each spectrum. If the software could provide a COSMIC score (Hoffmann *et al*., 2022), the candidate with the highest-ranking COSMIC score was selected. The corresponding SMILES and the compound classification provided by the CANOPUS (Dührkop *et al*., 2020) were extracted and stored for each spectrum and aggregated in a separate classification table. Compound classes provided by CANOPUS were analyzed at the CHEMONT level of subclasses and superclasses. The classes were aggregated and counted for each spectrum found in a sample and multiplied by the (log2-transformed) peak abundances of the corresponding MS1 precursors in the MS1-level peak table described above. The same procedure was accomplished for classes provided by CANOPUS regarding the Natural Product Classifier (NPC) (Gaudry *et al*., 2023).

Molecular descriptors were calculated for the SMILES provided by SIRIUS using the R package rCDK (Voicu *et al*., 2020) resulting in a data matrix with SMILES in rows and descriptors in columns. A data table was constructed corresponding to the feature table by performing a matrix operation of both tables. This data table was used for performing the data analyses.

Raw metabolite profiles and the annotated feature tables were deposited in the MetaboLights repository (study identifier MTBLS2239). Code to reproduce the results and generating the MAF for MetaboLights is available on GitHub (https://ipb-halle.github.io/iESTIMATE/doc/marchantiales.html).

### Data postprocessing

Morphometric measurements were transferred to a 2-dimensional matrix with measurements being binned, blanks padded and missing values imputed with zero. The resulting data matrix consisted of 160 rows and 23 columns. Traits with continuous numbers were subjected to additional mean, variance, skewness and kurtosis calculations to enrich the model selections resulting in a matrix with 160 rows and 91 columns.

Since the LC-MS was performed in positive and negative ionization modes, the molecular measurements were initially provided by two separate 2-dimensional feature tables with 48 rows and 17403 molecular features for negative mode and 29059 features for positive mode, respectively. Both tables were log-transformed to reach near normal distribution. To streamline statistical analyses, the distributions of both tables were inspected visually and tested using the Kullback-Leibler (KL) divergence for similarity. As the KL divergence was below 0.05, both tables were joined and, thus, positive and negative measurements did not require any separate treatments. Missing values of the resulting singular table (called “*feat_list*”) were imputed with zero resulting in a matrix with 48 rows representing the samples and 46462 columns representing the molecular features. To relate molecular features to annotated molecules and to improve subsequent analyses, a compound table (called “*comp_list*”) was created from the feature table that only contained those features that also had MS2-spectra associated with them. This resulted in a table with 48 rows and 5144 columns representing the molecules. From the compound table, a binary presence-absence table (called “*bina_list*”) was calculated. All values below 10% of the maximum feature intensity were set to zero and all values above to one. An additional binary table containing all unique features that were within a group of samples (species) but not the other (called “*uniq_list*”) was calculated by using the presence-absence table as a reference.

Molecular traits were estimated based on (1) computational annotation of small molecules using SIRIUS 5.7 (Dührkop *et al*., 2019b), (2) computational compound classification using CANOPUS as part of the SIRIUS software (Dührkop *et al*., 2019b, 2021; Hoffmann *et al*., 2022) and (3) predictions of molecular descriptors using the CDK framework based on the SMILES calculated by SIRIUS (Willighagen, 2017). 4314 out of 5144 spectra were annotated with class or putative structures.

The molecule annotations provided by SIRIUS were matched to the MS1 references in the compound table. A classification table (called “*class_list*”) was constructed by assigning and counting the most-specific classes compound class annotations for each annotated molecule per sample resulting in a classification table with 48 rows and 455 columns (representing 455 CHEMONT classes). Similarly, tables at the subclass level (called “*subclass_list*”) (140 columns), superclass level (called “*superclass_list*”) (16 columns), natural product class level (called “*npclass_list*”) (239 columns) and natural product pathway level (called “*nppathway_list*”) (7 columns) were generated.

Molecular descriptors were calculated for the SMILES provided by SIRIUS for every annotated molecule using rCDK version 3.8.1 (Voicu *et al*., 2020) resulting in a data matrix with 4314 SMILES in rows and 241 molecular descriptors in columns. Data tables were constructed corresponding to the compound table (called “*mdes_comp_list*”) and the presence-absence table (called “*mdes_bina_list*”) by performing a matrix operation of both respective tables resulting in tables with 48 rows (corresponding to the samples) and 241 columns (corresponding to the molecular descriptors).

### Data mining

For explorative analyses and to explore the spectrum of phenotypic and molecular traits, principal components analysis (PCA) was performed using the prcomp function in R. PCA scatterplots and biplots were accomplished using the autoplot function of the ggplot2 R package (version 3.5.1). In order to assess the explained variation of different study factors, variation partitioning was applied using the function varpart of the package vegan (version 2.6-6.1).

To evaluate the significance of morphometric traits, recursive and partitioning and regression trees (Gordon *et al*., 1984; Peters *et al*., 2018a) were calculated and model performance was determined using 10-fold cross-validation as implemented in the R packages caret (version 6.0-94) and rpart (version 4.1.23). The best tree was chosen from the model and visualized in R using a decision tree plotting method.

To evaluate the significance of molecular traits, variable selection was employed with Random Forest (RF) using the randomForest (version 4.7-1.1) and caret packages. A prediction model was trained using the train function from the caret package, and variable importance values were extracted from the model using the varImp function from the R package caret. Variables were selected (hence, were considered significant) when their variable importance (quantile threshold) was above 0.95. In order to visualize significant relationships of the selected variables at the different levels, heatmaps were generated from the selected variables using the gplots R package (version 3.1.3.1). To evaluate the performance of the fitted models, we additionally utilized 10-fold cross-validation (package mltest 1.0.1), and the Receiver Operating Characteristic (ROC) and PR (Precision and Recall) curves using the functions plot.roc and ci.se from the pROC 1.18.5 package and the function pr.curve from the PRROC 1.3.1 package were additionally constructed (Grau *et al*., 2015).

The R-squared of the fitted vs. the entire model and the area under curve (AUC) were calculated from the ROC, and the area under precision recall curve (AUC-PR) was determined from the PR curve.

Distance-based ReDundancy Analyses (dbRDA) were performed using the dbrda function of the package vegan in order to investigate relationships between pairs of different traits (i.e., between molecular traits and phenotypic traits, or between molecular traits and BET traits). BET traits refer to the Bryophytes of Europe Traits (BET) data set including a total of 55 biological traits such as those related to life-history, growth habit, sexual and vegetative reproduction, ecological traits such as indicator values, substrate and habitat, and bioclimatic variables based on calculated species ranges (Van Zuijlen *et al*., 2023). A Euclidean distance measure was used for the ordination. The dbRDA-model with the largest explained variance was chosen using forward variable selection and the ordistep function. The goodness of fit statistic (squared correlation coefficient) was determined for the selected variables from the dbRDA ordination model.

To conceptualize EMVs, we first utilized dbRDA to ordinate molecular traits with ecological traits. The dbRDA model reduced the number of BET traits having significant impact on molecular traits to a total of nine summarized by two largely orthogonal groups of traits that were either having a biological or a bioclimatic background. Significant molecular traits were then selected using RF (using variable importance larger than 0.95) as described above for the species and the nine significant BET traits resulting in ten lists of significant molecular traits that were correlated to ecological functioning. This procedure was accomplished at the levels of molecules, molecular family (NPC class) and molecular descriptors, respectively, and revealed a total of 2159 molecular traits (Table S1).

To identify EMVs, eligible molecular traits were narrowed down by filtering for only those variables that were shared in either one of the two groups of BET traits identified above. This resulted in a total of 468 candidate molecular traits. These 468 remaining molecular traits were then functionally annotated using secondary chemical information available in the libraries PubChem (Kim *et al*., 2019), LOTUS (Rutz *et al*., 2021), KEGG (Kanehisa *et al*., 2014), ChemFOnt (Wishart *et al*., 2023), and NPASS (Zeng *et al*., 2018). These libraries contain various types of chemical information that were manually curated for each molecular trait. If the exact molecular structure represented by the SMILES was not in the libraries, the search was broadened to structures using a Tanimoto similarity of 0.98. The contextual information obtained from the libraries was used to broadly functionally categorize them into the two groups of BET traits that were revealed by dbRDA to significantly constraining the molecular traits matrix: (1) biotic background (categories biological activity and metabolism and homeostasis), and (2) bioclimatic background (categories biosynthesis and plant growth and environmental adaptation). The percental partitioning of each of the categories was visualized by using a barplot using ggplot2.0. Finally, to select for candidate categories of EMVs, remaining molecular traits were grouped by their belonging to Natural Product Pathways and further manually cross-linked and curated based on their frequencies in the tables. This initially revealed eight categories, whereas Polyketides contained mostly amino acid-derived molecular traits and were joined with the category Amino acids and peptides, and fatty acids and carbohydrates were aggregated into the category Carbon metabolism as they were interdependent and PCA loadings were overlapping to a large degree.

Finally, this revealed molecular traits to be partitioned into six main categories of EMVs. These categories and subgroups of EMVs were collected into table 2 and their molecular function and proposed functioning at ecosystem scales was obtained with literature research. To visualize EMVs and their relevance in the entire dataset, a circular heatmap of molecular traits belonging to the six categories of EMVs was accomplished using the R packages circlize 0.4.17 and ComplexHeatmap 2.24.1. To show the relevance of EMVs, an ordination using dbRDA of molecular traits belonging to the six categories of EMVs and the nine BET traits identified above and species identity was carried out and visualized using aes (ggplot) and envfit functions of ggplot2 3.5.2 and vegan 2.7-1 R packages.

**Table 2.**
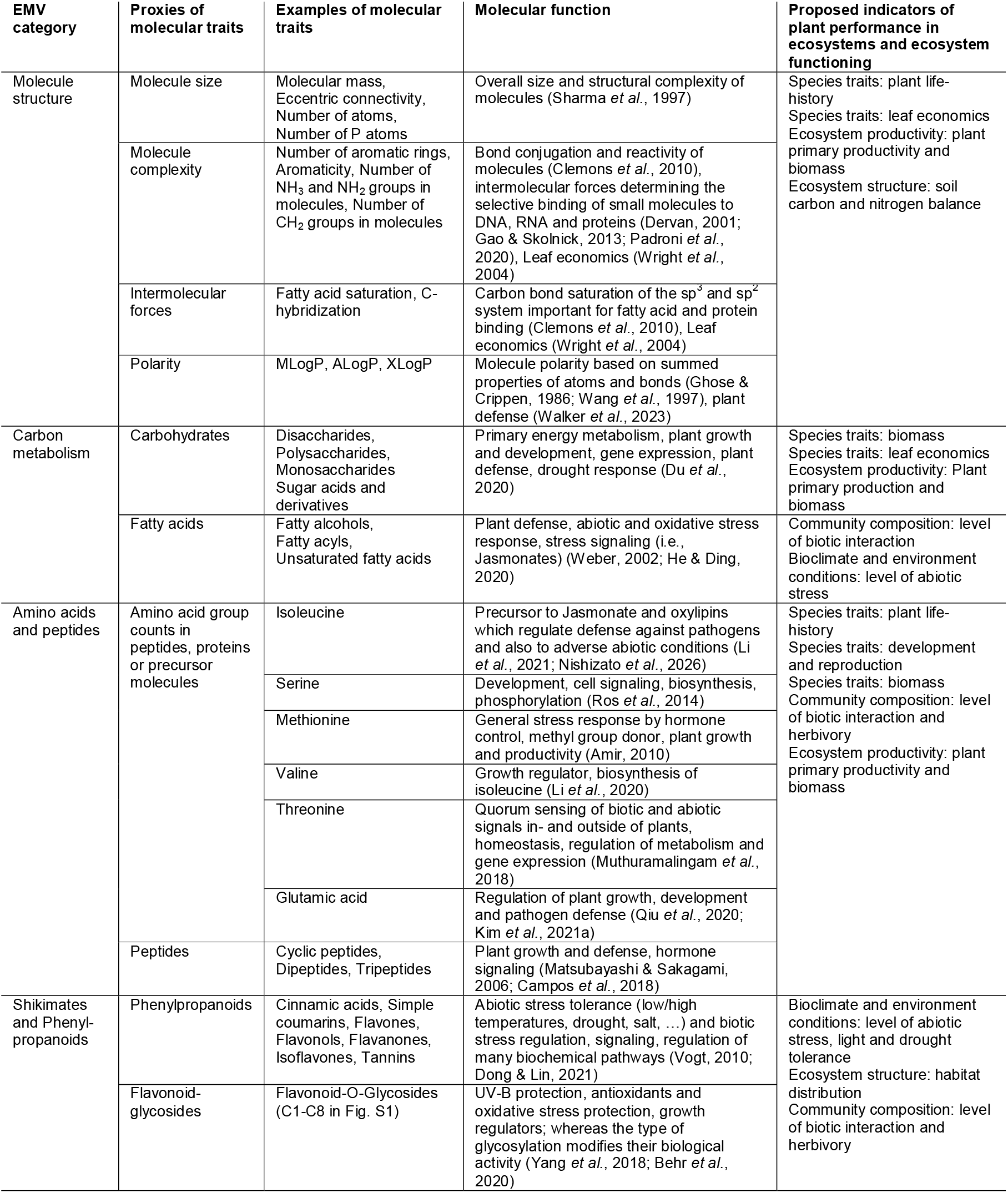
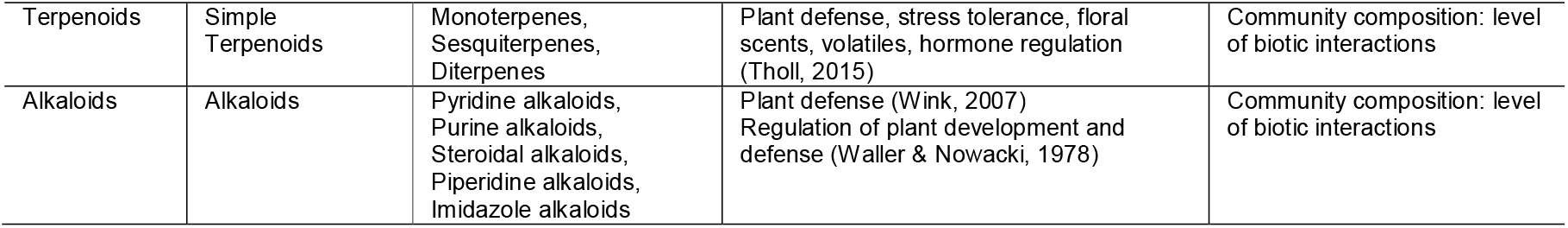
Six categories of EMVs. They follow a hierarchy. Each category contains proxies of molecular that group together several molecular traits. Examples are given in the third column. The column molecular function describes the mechanisms at the cellular level. The last column proposes their role as indicators of plant performance in ecosystems and general ecosystem functioning. To propose indicators, we followed the terminology of essential biodiversity variables listed at the https://geobon.org website.

## Results

The 23 acquired morphometric traits were analyzed using a recursive partitioning and regression tree. The rtree model was able to differentiate 12 of the 13 species based on morphometric traits (Fig. 2a), with *Riccia beyrichiana* not included in the model due to low model confidence. The rtree model selected five morphometric traits as highly indicative for species identity (Fig. 2a). Interestingly, the rtree model differentiated the morphologically similar *Riccia gothica* and *R. subbifurca* by thallus width in the cross-section, which is in accordance with the *species novo* description and recent taxonomic investigations in the genus *Riccia* (Damsholt & Hallingbäck, 1987; Pöltl *et al*., 2025).

**Figure 2.**
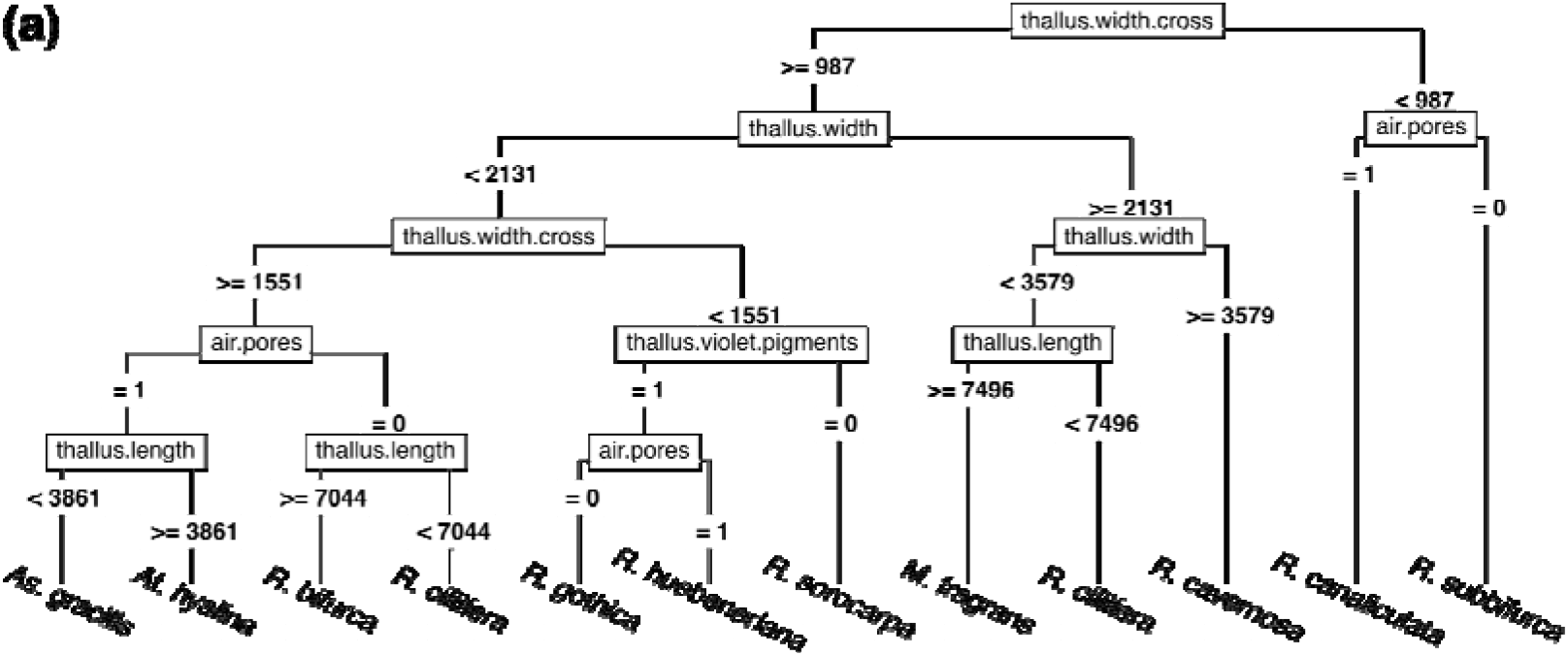
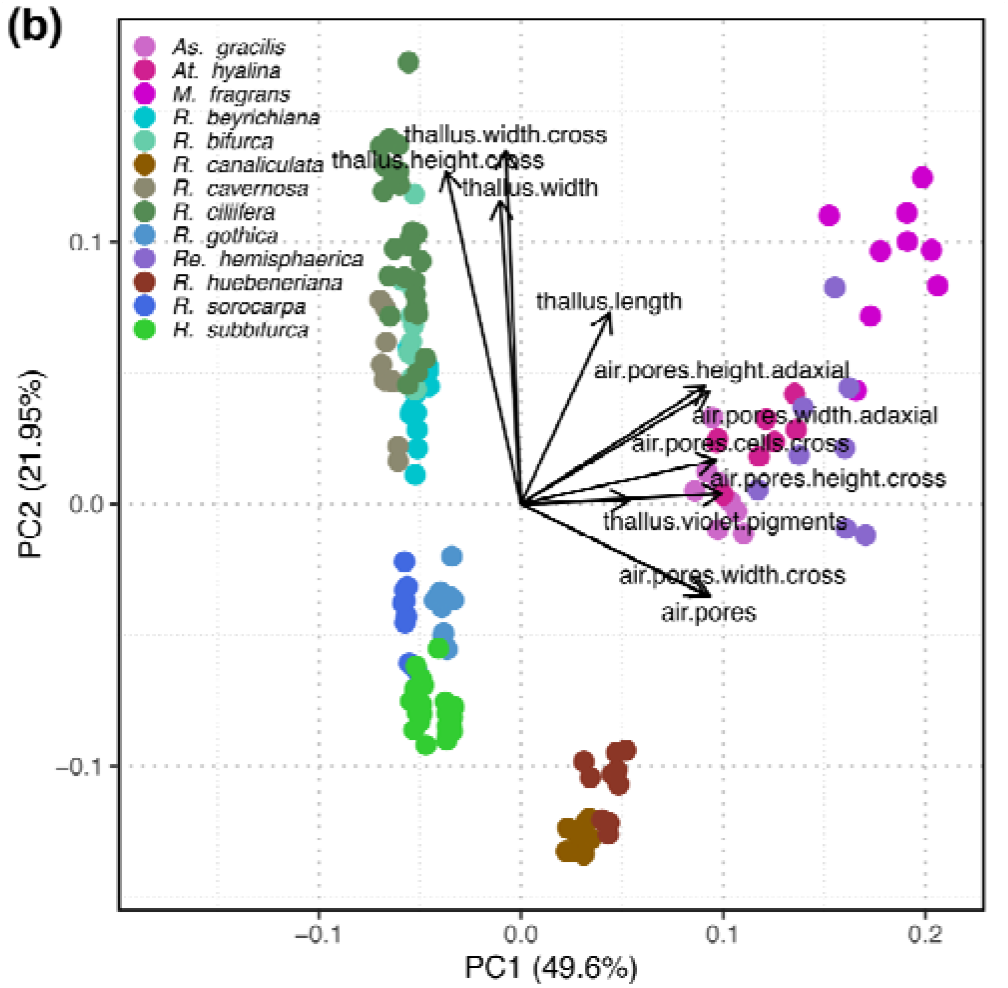
**(a)** Recursive partitioning and regression tree of morphometric traits acquired for the 16 different samples. For full species names, see table 1. R^2^=0.654, multi-class accuracy=0.695. **(b)** PCA biplot of the selected morphometric traits by the rtree model. Explained variance on PC1=49.60% and PC2=21.95%.

To get an overview on which components dominate the dimensions of the classical trait spectrum of the investigated complex-thallose liverworts, a PCA biplot was performed of the selected morphometric traits by the rtree model (Fig. 2b). The selected traits corresponded mainly to the size, structure and thickness of air pores at the first PC axis and to width/length ratios of the thallus at the second PC axis (Fig. 2b).

### The structure of the molecular trait spectrum

We performed ordinations of the molecular traits grouped by the main molecular pathways as defined by the Natural Product Classifier (NPC) (Gaudry *et al*., 2023) with life-history traits obtained from the BET data set (Van Zuijlen *et al*., 2023) (Fig. 3a) as well as with the morphometric traits obtained in this study (Fig. 3b). Whereas the composition and diversity of molecular traits explained more variance (PC1: 49.9%) than life-history traits (PC2: 18.9%), they were also largely orthogonal with the exception of generation length (genl), life-strategy (lstrat) and plant size (Fig. 3a). The ordination of molecular and morphometric traits showed molecular traits to be largely orthogonal as well, but explaining less variance (PC2: 24.0% vs. PC1: 35.2% variance) (Fig. 3b).

**Figure 3.**
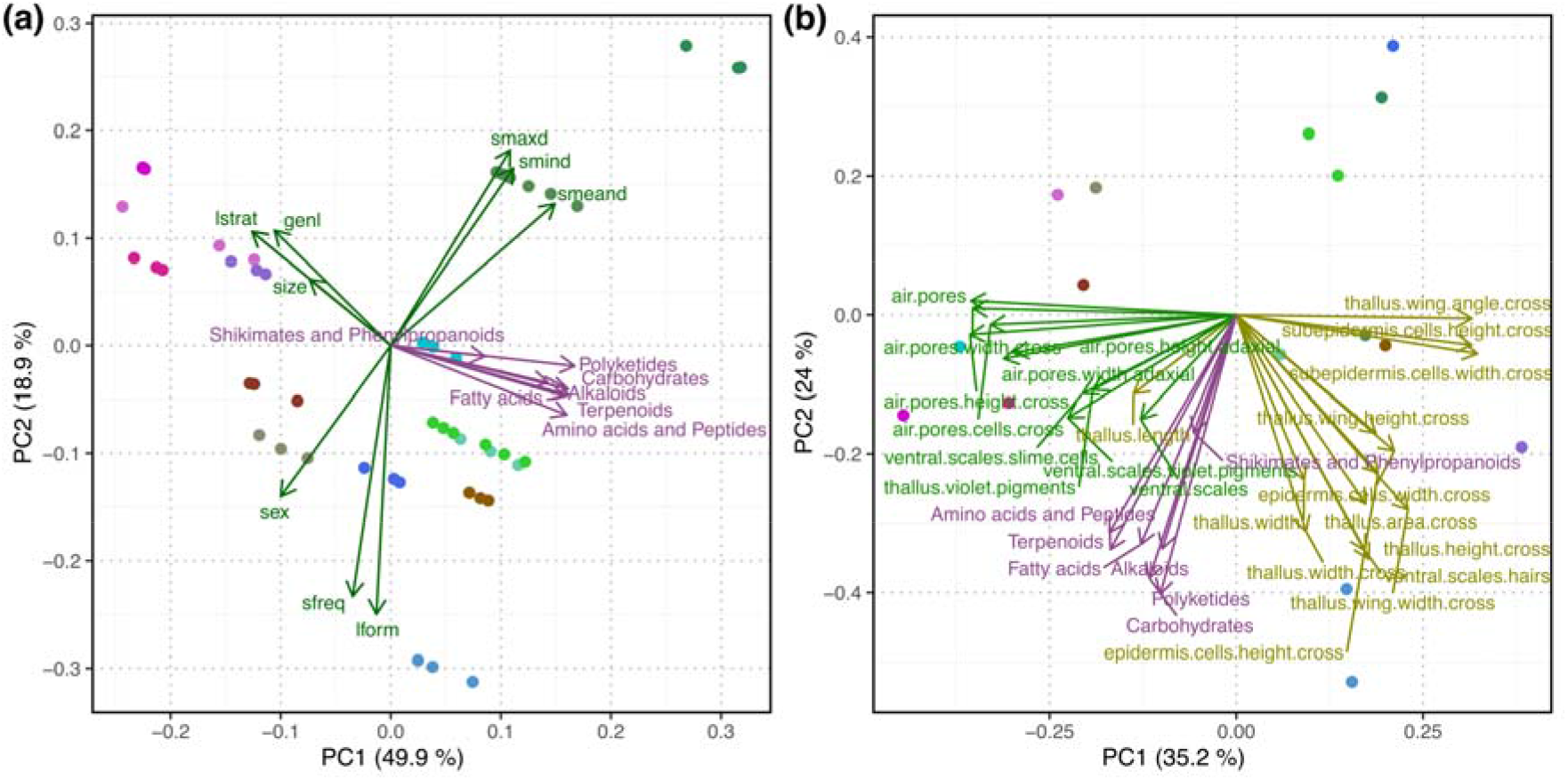
**(a)** PCA biplot of molecular pathways (purple color) and BET traits (green color) combined. Selected BET traits: lstrat=life strategy, genl=generation length, smaxd=maximum spore diameter, smind=minimum spore diameter, smeand=mean spore diameter, sex=mating type/sexual condition, sfreq=sporophyte frequency, lform=life form. Explained variance on PC1=49.9% and on PC2=18.9%. **(b)** PCA biplot of molecular pathways (purple color) and obtained morphometric traits (thallus and cell measurements in yellow color, air pores and ventral scale structures in green color). Explained variance on PC1=35.2% and on PC2=24.0%.

### Essential Molecular Variables (EMVs)

To identify major categories of EMVs, we first employed ordination using dbRDA of the molecular traits with the 55 bryophyte traits of BET (Van Zuijlen *et al*., 2023). dbRDA reduced the number of BET traits having significant impact on molecular constitution to a total of nine summarized by two largely orthogonal groups that were either having a (1) physiological and biotic background or related to (2) bioclimatic conditions (Fig. 4a). These two groups also defined the first two principal components structuring most of the molecular form and function (37.18% of the total variation, see Fig. 4a).

**Figure 4.**
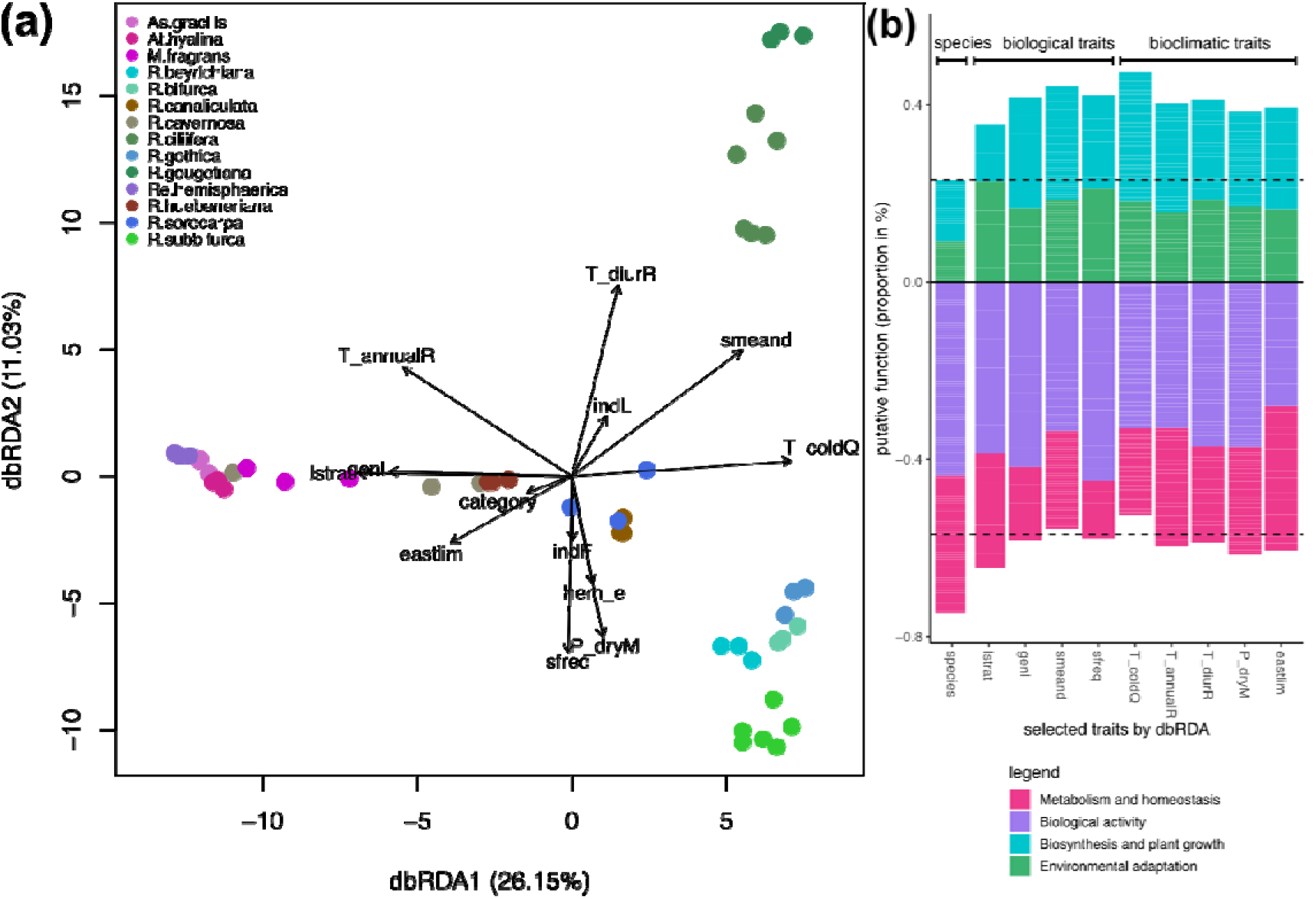
**(a)** dbRDA of molecular traits and BET traits. Species samples are shown as colored dots and traits are overlayed using arrows. The longer the arrow the higher the significance and the closer the samples to the arrow the closer the relationship. Explained variance on axis 1: 26.2% and on axis 2: 11.0%. For full species names, see table 1. **(b)** Barplot showing the proportions (%) of putative functions of molecular traits to the significant BET traits revealed by dbRDA. Putative functions are grouped by (1) intrinsic bio-molecular processes within plants such as metabolism, homeostasis and biotic background (red- and purple-colored bars), and (2) indication of extrinsic biogeochemical processes such as biosynthesis, plant growth and environmental adaptation (green and blue colored bars). The first column represents functions related to species identity. The next columns show functional annotations of molecules into (1) biological traits composed of life strategy (lstrat), generation length (genl), mean spore diameter (smeand), and sporophyte frequency (sfreq), and (2) bioclimatic traits composed of mean daily mean air temperatures of the coldest quarter (T_coldQ), annual range of air temperature (T_annualR), eastern limit category (eastlim), precipitation amount of the driest month (P_dryM), and mean diurnal air temperature range (T_diurR).

Next, molecular traits having significant relation to the nine selected BET factors were determined using Random Forest (RF), individually at the levels of molecules, molecular family (natural product class) and molecular descriptors, revealing 2159 of potential candidates of molecular traits. To narrow down the list of candidates, a data reduction strategy was accomplished by selecting only those molecular traits with RF variable importance larger than 0.95 and by limiting to only those that were shared in either one of the two orthogonal groups of traits identified above. This revealed a total of 468 candidate molecular traits (Table S1).

To identify EMVs, the 468 remaining molecular traits were grouped according to molecular family and functionally annotated using the chemical libraries PubChem (Kim *et al*., 2019), LOTUS (Rutz *et al*., 2021), KEGG (Kanehisa *et al*., 2014), ChemFOnt (Wishart *et al*., 2023), NPASS (Zeng *et al*., 2018) and with additional literature research. The information for each molecular trait was manually curated and aggregated, and the traits were grouped by molecular family and for their putative function in relation to (1) biotic interactions and physiological background explaining the molecular function regarding the biological activity and intrinsic bio- molecular processes within plants such as metabolism, homeostasis; and (2) their putative bioclimatic functioning regarding extrinsic biogeochemical processes such as biosynthesis and plant growth, or environmental adaptation (Fig. 4b). The traits were then grouped by molecular family, aggregated according to their putative functioning, sorted by their variable importance and manually cross-linked.

This revealed that molecular traits were arranged in six categories of EMVs summarized in Table 2. They comprise (1) molecule structure (size, mass, complexity, arrangement, aromaticity, connectivity and polarity of atoms in molecules), (2) molecules involved in the carbon metabolism (carbohydrates such as mono-, di- and polysaccharides, sugar acids, and fatty acids such as fatty alcohols, fatty acyls and unsaturated fatty acids), (3) amino acids and peptides (counts of amino acid groups in molecules and cyclic, di- and tripeptides), (4) shikimates and phenylpropanoids (general phenylpropanoids such as cinnamic acids, coumarins, (iso)flavones, flavonols, flavanones and tannins and specific flavonoid-glycosides), (5) terpenoids (mostly simple mono-, sesqui- and diterpenoids), and (6) alkaloids (pyridine, purine, piperidine, imidazole and steroidal alkaloids) (Table 2).

To validate and visualize the functional profile of EMVs, a circular heatmap was employed combined with dendrograms and correlation coefficients indicating the similarity of traits within the EMV categories (outer parts in Fig. 5). In the center of the plot in Fig. 5, a PCA of EMV categories and the selected bioclimatic factors is showing the distinctive functional profiles of the EMVs. The first PC axis is largely corresponding to species and their life strategies (37.7%), whereas bioclimatic factors are largely corresponding to the second PC axis (12.8%). The EMVs flavonoid-glycosides and phenylpropanoids show the strongest relationship with bioclimatic factors (light and dark green ellipses in the center of Fig. 5), whereas the EMVs terpenoids, alkaloids and carbon metabolism (purple, orange and red ellipses in the center of Fig. 5) additionally capture biotic factors. These functional profiles are confirming the generalized plant performances as shown in the last column in Table 2.

**Figure 5.**
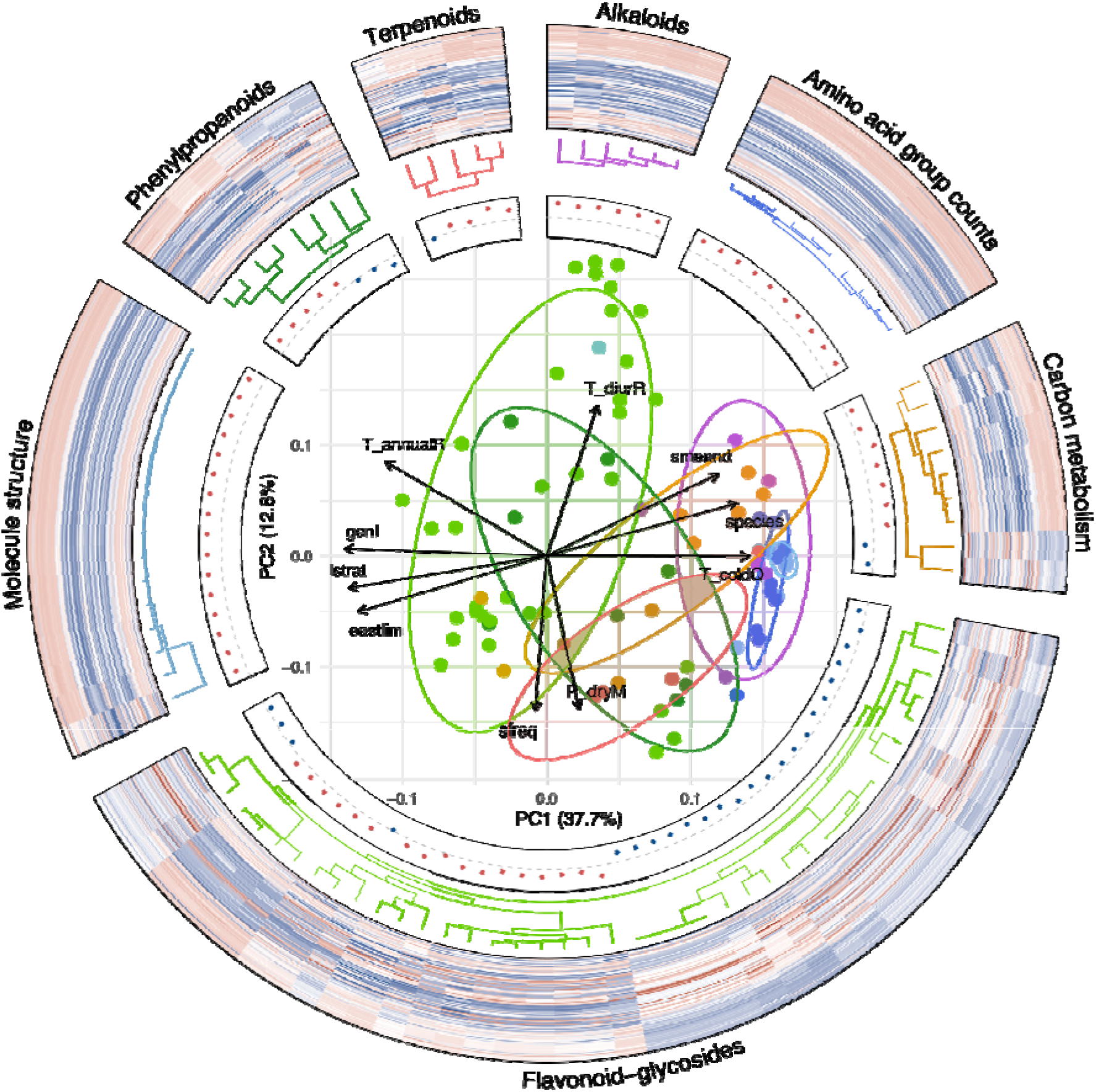
Plot showing the distribution and functional profiles of EMVs. The heatmaps show the distribution of molecular traits among the investigated species. A red color indicates a positive relative abundance while a blue color indicates negative relative abundance. A white color indicates no abundance. On the inside, dendrograms show the relationships of traits within the EMVs. Further to the inside, correlation coefficients of traits within EMVs are shown, whereas a red color indicates a positive and a blue color a negative correlation. In the center of the plot, a PCA plot of EMV categories and the selected bioclimatic factors is showing the scores of the EMV constituents, whereas ellipses indicate 80% of confidence levels. Arrows are showing the selected nine bioclimatic factors and their relation to the EMVs. The closer a score to the arrow the larger the correlation. The meaning of the bioclimatic factors is explained in Fig. 4. Light green: Flavonoid-glycosides, dark green: Phenylpropanoids, Red: Terpenoids, Orange: Carbon metabolism, Dark blue: Amino acid group counts, Light blue: Molecule structure, Purple: Alkaloids.

## Discussion

### The structure of the classical trait spectrum

Despite the large variation of form and function in liverworts, it has been rarely investigated how traits vary among taxonomical groups and whether few essential traits combinations (traits syndromes) constrain the multi-dimensional trait matrix (Waite & Sack, 2010; Coe *et al*., 2024). We found the classical spectrum to be mainly structured by the size, structure and thickness of air pores and width/length ratios of the thallus which also separated the investigated complex-thallose liverwort species (Fig. 2a,b). The species-specific variations in the surface leaf-area reflect differences in light capture, whereas the variations in thallus thickness and differences in the structure of air pores represent trade-offs between carbon gain and water transpiration (Grau-Andrés *et al*., 2022). Thallus thickness and width/length ratios (and hence thallus volume) are strongly correlated to the leaf mass per area (LMA) trait that is one of the main components of leaf economics of vascular plants (Stanton & Coe, 2021). As many of the traits on the first axis (structure and sizes of air pores) are related to water balance, they underpin the importance of desiccation tolerances in bryophytes and the pronounced trade-offs between reproduction, growth and water availability in biocrusts (Seppelt *et al*., 2016).

Taken together, in the investigated liverworts we find a phenotypic trait spectrum similar to that of vascular plants as the first dimensions were dominated by classical traits related to the size of whole plants (thallus lengths and widths) and traits related to the leaf economics (thallus thickness, air pores structures). This confirms that fundamental relationships of form and function described for vascular plants also apply to liverworts (Violle *et al*., 2012; Lamanna *et al*., 2014; Díaz *et al*., 2016).

However, traits related to air pores size and structure highlight the importance of water relations and desiccation tolerances, which makes the economics spectrum of bryophytes differently composed to that vascular plants (Stanton & Coe, 2021).

### The structure of the molecular trait spectrum

To assess how molecular traits integrate with the classical trait spectrum, we performed ordinations of the molecular traits with life-history traits obtained from the BET data set (Van Zuijlen *et al*., 2023) (Fig. 3a), as well as with the morphometric traits obtained in this study (Fig. 3b). Molecular traits were largely orthogonal to classical traits. Thus, they explain additional axes of functional specialization also in liverworts (Walker *et al*., 2022). In addition, we found molecular traits to be mainly related to mechanistic adaptation processes to bioclimatic changes and regulatory adaptations within plants and life-history and conclude that the fundamental relationships of form and function also apply at the molecular level (Violle *et al*., 2012; Walker *et al*., 2023). However, as the investigated liverworts are mainly characterized by desiccation tolerances in contrast to tracheophytes, the spectrum of form and function is likely to be explained by different sets of individual traits (Coe *et al*., 2024). We propose to investigate the molecular variation of traits such as dynamic water content, specific thallus mass, thallus mass per area, thallus N content, or organizational scale (growth in colonies, mats, cushions) (Stanton & Coe, 2021) to further substantiate the fundamental relationships of form and function in liverworts.

### Essential Molecular Variables (EMVs)

Molecular traits are typically used to explain mechanisms within plants and have become an indispensable part of molecular biology and biochemistry (Pichersky & Lewinsohn, 2011; Weng *et al*., 2021). To make use of molecular traits in ecology, we first identify patterns of molecular traits resulting from changes at ecological scales, categorizing them and scaling up selected categories of molecular traits again to spatiotemporal coarser ecological scales to validate the relationships (Peters *et al*., 2018b).

We found the molecular trait spectrum to be highly structured and consistent among species and across taxonomic groups. They originate from distinct evolutionary histories of species and types of biocrusts. Proxies of molecular traits were arranged in six major categories of EMVs summarized in Table 2. They comprise (1) molecule structure, (2) molecules involved in the carbon metabolism, (3) amino acids and peptides, (4) shikimates and phenylpropanoids, (5) terpenoids, and (6) alkaloids.

Each of the EMVs is showing a distinctive functional profile capturing proxies of molecular traits either relating to (1) biotic interactions and physiological background, or to (2) bioclimatic functioning relating to abiotic environmental conditions and biosynthesis and plant growth (Table 2, Fig. 4b, Fig. 5). This functional information directly relates to their performance in ecosystems. The extracted EMVs are thus also likely indicative for ecosystem functioning which mean they inform ecology on the molecular mechanisms underlying biotic interactions or bioclimatic changes and capture major dimensions complementary to one another (Fig. 5).

The EMV category molecule structure comprises molecular traits that relate structural properties of molecules such as molecule size, structural complexity, polarity and intermolecular forces to be indicative for leaf economics, plant primary productivity and life-history in ecosystems. They link molecular biosynthesis, the primary energy metabolism and gene expression to impact the ecosystem’s primary production via changing the overall plant biomass, soil carbon and nitrogen balance. In particular, aromaticity, C-hybridization and fatty acid saturation descriptors are closely related to the composition and abundance of various saccharides and (unsaturated) fatty acids summarized by the EMV carbon metabolism. Molecular mass, number and count of atoms, CH_2_- and NH_3_-groups in molecules are indicative for resource acquisition and carbon metabolism; intermolecular forces like c-hybridization ratios are important for fatty acids saturation; polarity, double-double C-bonds part of benzenes and terpenoids, and number of rings affect bond conjugation, reactivity of molecules and are indicative for plant defense (Ghose & Crippen, 1986; Wang *et al*., 1997). While molecular traits related to molecule structure, amino acid and carbon metabolism have been found to vary predictably among taxonomic groups before (Walker *et al*., 2023), our analysis additionally revealed their ecological functioning, hence, that they predominantly predict life-histories and indicate general plant strategies that regulate trade-offs between growth and reproduction in different ecosystems.

In the EMV category amino acids and peptides we identified isoleucine, serine, methionine, valine, threonine and glutamic amino acid group counts in molecules and cyclic, di- and tripeptides to be indicators of development, reproduction and growth affecting ecosystem productivity via trade-offs in the plants’ life-history and underlying pathogen and herbivory responses. At the molecular level, they play important roles either as intermediates in biochemical pathways or as plant hormones regulating cell signaling, development, phosphorylation, biosynthesis, or various defense mechanisms. For example, arginine, or aspartic acid act as nitrogen reserve, recycling and precursors to e.g. NAD (Han *et al*., 2021), isoleucine and valine act as precursors to Jasmonate significantly regulating plant defense and plant stress responses to adverse abiotic conditions (Li *et al*., 2021), or methionine involved in stress response, plant growth and productivity (Amir, 2010). The upregulation of di- and tripeptides suggests that liverworts invest a large degree of molecular resources into biosynthesis under stressful conditions similar as with vascular plants (Volpes *et al*., 2023). Cyclic peptides have been described to have strong antimicrobial properties suggesting that plants also structure the community through biosynthesis of these molecules (Campos *et al*., 2018).

The shikimate and phenylpropanoid pathways are central to plants and are long known to play key roles with stress induction (Dixon & Paiva, 1995; Vogt, 2010). In this EMV category we found cinnamic acids, simple coumarins, flavones, flavonols, (iso)flavanones, tannins and especially flavonoid-glycosides to be indicative for different kinds of abiotic stress (light, drought, temperature, pH) and moreover also characterizing the ecosystem structure in different kinds of environments. In particular, glycosides and flavonols are attached to each other via the flavonoid 3-O- glucosyltransferase (UFGT). Despite that the detected flavonoid-glycosides are commonly produced in all land plants, our results clearly show that the composition and abundance is indicative for differences in bioclimatic conditions, long- and short-term abiotic habitat fluctuations, or disturbance regimes that indicate the ecosystem structure and habitat distribution at coarse ecosystem scales (Table 2). In literature, flavonoid-glycosides have been described to carry out multiple functions regarding oxidative stress and changes in light, drought or temperature regimes (Le Roy *et al*., 2016). A large diversity of flavonoid-glycosides is giving plants evolutionary advantages in different habitats while representing immediate and flexible molecular responses to short-term environmental changes (Kulshrestha *et al*., 2022). This confirms earlier findings of molecular diversification due to different bioclimatic conditions (Markham & J. Porter, 1975; Markham *et al*., 1976; Davies *et al*., 2020). As shown for flavonoids, molecules are likely to carry out multiple cellular functions due to different (post-)transcriptional regulations, including protein confirmations and glycosylation, as well as in ecology where molecules likely serve multiple functions which increases resilience to stress and overall fitness, but exacerbating the functional and mechanistic characterization (Zanzoni *et al*., 2019; Sack & Buckley, 2020; Horn *et al*., 2021; Nascimento & Tattini, 2022). The high diversity in flavonoid-glycosides can be explained by the fact that plants modify their biological activity by choosing different types of glycosylation depending on environmental conditions (Yang et al. 2018; Behr et al. 2020).

The two EMV categories terpenoids and alkaloids contain proxies of molecular traits that regulate stress tolerance, play roles in hormone production, as volatiles, in plant development and defense mechanisms. Plants produce specific kinds of terpenoids and alkaloids as a response to changes in biotic conditions such as production of floral scents to attract pollinators, mobilization of beneficial microbial and fungal partners, or bioactive agents to repel pathogens or herbivores (Cheng *et al*., 2007; Pichersky & Raguso, 2018). The composition and diversity of molecules and molecular classes within these two EMVs are characteristic for the levels and kinds of interactions with other organisms and directly link to the community composition (Table 2). We found a large diversity of mono-, sesqui- and diterpenes and pyridine, purine, piperidine and imidazole alkaloids that were described to exhibit antimicrobial, antifungal and cytotoxic activities suppressing the activity of microbials, insects and other invertebrates and, thus, structuring (microbial) communities in the ecosystem (Asakawa *et al*., 2013; Eldridge *et al*., 2023). Variations in abundance and kinds of steroidal alkaloids have also been described to regulate plant development and plant defense mechanisms, further underlining their fundamental role in plants responses to biotic changes (Wink, 2007).

Finally, using our example of liverworts in biocrusts we want to provide examples on how EMVs can generate new hypotheses and improve our mechanistic understanding of plant performance in ecosystems. As temperatures and the frequency of precipitation were reported to be the main drivers of composition and biodiversity of crusts (Navas Romero *et al*., 2020) and climatic seasonality has been shown to limit soil functions and nitrogen availability to bryophytes (Eldridge *et al*., 2023), consequently increasing extinction risks (Van Zuijlen *et al*., 2024), the investigated liverworts may be vulnerable to extreme weather conditions predicted to increase with global change, especially when extremes like absence of rain or temperatures well-above the optima of the species occur during summer. This was shown for *Syntrichia caninervis* and other mosses and selected bioclimatic traits in combination with narrow niches greatly increase extinction risks (Gignac, 2001; Reed *et al*., 2012; Henriques *et al*., 2016; Hodgetts, 2019; Van Zuijlen *et al*., 2024). While the investigated annual *Riccia* species likely either produce incomplete thalli lacking sporophytes in autumn, thalli of bi- or perennial thallose liverworts may take damage and growth and sporophyte formation, which occurs during spring and summer, will be severely reduced (Clausen, 1964; Furness & Grime, 1982; Seppelt *et al*., 2016). In this regard, the monoecy in the investigated *Riccia* species (except *R. ciliifera*) can already be viewed as adaptation to sites with high bioclimatic variability greatly depending on water availability during sporophyte formation (Coe *et al*., 2024; Levins *et al*., 2025). With a phenotypic trait-based approach it was shown that bryophytes are functionally sensitive to small-scale environmental variability and that traits related to water balance have pronounced effects (Deilmann *et al*., 2025). We also found highly diverse fatty acyls that are part of the cutin and waxy layer of thallose liverworts. They regulate water relations and play important roles in the desiccation tolerances of bryophytes (Davies *et al*., 2020; Nascimento & Tattini, 2022; Munoz *et al*., 2024). In bryophytes, they further play roles in oxylipin biosynthesis and Jasmonate pathways (Nishizato *et al*., 2026).

We hypothesize that extreme events could destabilize biocrust communities by removing the dominant bryophytes from the species pool disturbing the functional profile of biocrusts, leaving them exposed and further vulnerable (Steven *et al*., 2015) and likely converting to an earlier successional stage (Ferrenberg *et al*., 2015). As a result, biodiversity in biocrusts will likely have higher turnover, either resulting in the loss of highly specialized species, or the replacement with few generalist species. The presence of the many molecules annotated to have biological activity also suggests that associated microbials and other organisms in soil crusts might additionally become at risk (Liang *et al*., 2020).

A high molecular diversity is characteristic for environments with high levels of abiotic stress and increases levels of facilitation among species in the community (stress-dominance hypothesis) (Swenson & Enquist, 2007). As thallose liverworts can exude small bioactive metabolites through their entire surface (Vicherová *et al*., 2020), they likely structure crust communities by adding beneficial or removing pathogenic microbial species from the species pool similarly to peat mosses (Vicherová *et al*., 2017; Sytiuk *et al*., 2022). It has been reported that thallose liverworts exchange specialized metabolites and form close environment-specific associations with nitrogen-fixing cyanobacteria (Rousk *et al*., 2013), actinobacteria (Delgado-Baquerizo *et al*., 2018), betaproteobacteria, methylobacteria and fungi (Wicaksono *et al*., 2023; Cheng *et al*., 2024). These low-abundant taxa have been considered to be key in the stability of crusts as they are involved in biogeochemical cycles, increase resilience against physical and bioclimatic disturbance and support enzymatic activity in soils (Banerjee *et al*., 2018; Delgado-Baquerizo *et al*., 2018; Eldridge *et al*., 2023). It is still unresolved whether or how essential microbials affect liverwort biology and may also drive their evolution through the production and timing of reproductive organs thereby facilitating their own spreading.

In summary, the high diversity of bioactive molecules and the spectrum of significant molecular descriptors confirms our earlier hypothesis that liverworts predominantly interact with their biotic and abiotic environment through specialized metabolites (Peters *et al*., 2022). Using EMVs to disentangle the molecular interactions and responses of liverworts in biocrusts further helps to understand how they regulate microbial communities, protect soils and to mitigate the effects of global change (Delgado-Baquerizo *et al*., 2018; Eldridge *et al*., 2023).

In conclusion, we found a phenotypic trait spectrum similar to that of vascular plants as the first dimensions were dominated by classical traits related to the size of whole plants (thallus lengths and widths) and traits related to the leaf economics (thallus thickness, air pores structures). However, economics traits in the investigated liverworts were composed of air pores structural and size-related traits that are different to those in vascular plants highlighting the importance of water relations and desiccation tolerances in bryophytes (Stanton & Coe, 2021). The molecular trait spectrum was highly structured by six categories of EMVs comprising (1) molecule structure, (2) molecules involved in the carbon metabolism, (3) amino acids and peptides, (4) molecules produced in the central shikimates and phenylpropanoid pathways, (5) terpenoids, and (6) alkaloids. EMVs provide detailed insights into mechanistic adaptation processes within the investigated liverworts but are also highly indicative for liverwort performance in ecosystems. They are linking molecular processes mechanistically to climate seasonality, environmental disturbance, biotic interactions, species turnover and life-history. Whether they are universal for describing terrestrial plant performance at both, molecular and ecosystem scales remain to be tested on vascular plants. By “zooming in” and, thus, identifying the individual constituents within the categories of EMVs at cellular levels, molecular mechanisms can be elicited (Peters *et al*., 2018b). A detailed investigation in other kinds of plants can generate new hypotheses and allow for follow-up experiments. The Supplemental Material contains some further molecular investigations of thallose liverworts that are beyond the scope of this paper but can serve as examples. Furthermore, by “zooming out” the mechanisms on how EMVs and individual molecular traits impact or are impacted by life-histories, bioclimate, or ecosystem stability can be revealed, uncovering more and broader functioning at ecosystem scales that requires abstraction of the detailed molecular data (Peters *et al*., 2018b). Substantiating EMVs and integrating them into an ecological context allows for the discovery of new mechanisms in, both, functional ecology and molecular biology. As so far molecules only provided limited insights into ecological functioning, investigation of EMVs and identification of proxies of molecular traits can open a new chapter bridging chemical with functional ecology by offering a new tool set for unravelling molecular mechanisms underlying plant performance in ecosystems and, thus, enhancing the explanatory power of the functional trait concept (Fraenkel, 1959; Hartmann, 2008; Weng *et al*., 2021; Walker *et al*., 2023).

## Acknowledgments

KP acknowledges the support of iDiv (funded by the German Research Foundation, DFG-FZT 118, 202548816). Further, we like to thank the Leibniz Foundation for supporting this study. We would also like to thank Jörg Ziegler for assistance in the lab and Helge Bruelheide and Christoph Rosche for valuable corrections to the manuscript. Species identities were kindly validated by Tomas Hallingbäck and Galin Gospodinov.

## Author contributions

KP conceptualized and performed the entire study and wrote the manuscript. SN acquired funding and reviewed and edited the final draft. NvD reviewed and edited the final draft. All authors contributed to the final version of the manuscript.

## Data availability

- Bioimaging data: The raw images (Canon CR3-format), the pre-processed images (16-bit TIFF-format) and the contextual metadata were deposited to BioStudies under the identifier S-BIAD824 (https://www.ebi.ac.uk/biostudies/studies/S-BIAD824). The data record consists of a total of 102’375 individual raw image files partitioned into 16 samples. The entire data record has a total size of approx. 8 TB. The pre- processed and processed images along with metadata were deposited to the Image Data Resource (IDR) repository under the identifier idr0157 (https://idr.openmicroscopy.org/search/?query=Name:157). The data record consists of a total of 1127 pre-processed and 157 fully processed imaged files. The data record has a total size of approx. 12 TB. Fully segmented images showing the species are available in Zenodo (https://doi.org/10.5281/zenodo.10683968).
- Molecular traits data: Raw metabolite profiles (zipped vendor data and in mzML format) have been made available in MetaboLights under the study identifier MTBLS2239 (https://www.ebi.ac.uk/metabolights/editor/MTBLS2239). The dataset includes 48 metabolite profiles in positive and negative modes, QC and blank profiles, metabolite feature tables (MAF) and metadata.
- Extracted tables and code used for analysis were deposited to Zenodo (https://doi.org/10.5281/zenodo.21137547). The dataset includes the RData containing all data objects and the R vignette containing code to reproduce to plots used in this study.
- Sequencing data were deposited to the European Nucleotide Archive (ENA) and are available under the study identifier ERP155252 (accession PRJEB70317) (https://www.ebi.ac.uk/ena/browser/view/PRJEB70317). Raw reads are available under the sample identifiers SAMEA114863468- SAMEA114863483.
- Metadata to voucher specimens are available at the virtual herbarium JACQ with the following identifiers: JE4010742, JE4010741, JE4010739, JE4010740, JE4010749, JE4010752, JE4010747, JE4010743, JE4010753, JE4010748, JE4010746, JE4010754, JE4010745, JE4010744, JE4010750, JE4010751.

